# Stem cell transcriptional profiles from mouse subspecies reveal *cis*-regulatory evolution at translation genes

**DOI:** 10.1101/2023.07.18.549406

**Authors:** Noah M. Simon, Yujin Kim, Diana M. Bautista, James R. Dutton, Rachel B. Brem

## Abstract

A key goal of evolutionary genomics is to harness molecular data to draw inferences about selective forces that have acted on genomes. The field progresses in large part through the development of advanced molecular-evolution analysis methods. Here we explored the intersection between classical sequence-based tests for selection and an empirical expression-based approach, using stem cells from *Mus musculus* subspecies as a model. Using a test of directional, *cis*-regulatory evolution across genes in pathways, we discovered a unique program of induction of translation genes in stem cells of the Southeast Asian mouse *M. m. castaneus* relative to its sister taxa. We then mined population-genomic sequences to pursue underlying regulatory mechanisms for this expression divergence, finding robust evidence for alleles unique to *M. m. castaneus* at the upstream regions of the translation genes. We interpret our data under a model of changes in lineage-specific pressures across *Mus musculus* in stem cells with high translational capacity. Our findings underscore the rigor of integrating expression and sequence-based methods to generate hypotheses about evolutionary events from long ago.

## Introduction

The central objective of molecular-evolution research is to draw inferences about changing selective pressures between lineages, based on clues from omics data. Since its inception, the field has relied in large part on model-fitting methods using DNA sequence (Kreitman 2000) and gene expression (Price et al. 2022). Complementing the latter, empirical genome ranking/bootstrapping approaches have emerged in the more recent literature (Ferguson and Chang 2020), for which methods development and refinement remain an active area of research (Berg et al. 2019; Sohail et al. 2019; Johri et al. 2020; Price et al. 2022).

One clear-cut empirical molecular-evolution strategy (Bullard et al. 2010; Fraser et al. 2010; Fraser et al. 2011; Martin et al. 2012; York et al. 2018; Gokhman et al. 2021) takes as input measurements of *cis*-regulatory variation from expression profiles. The test identifies cases in which, among the unlinked genes of a pathway subject to *cis*-regulatory change, alleles in one taxon tend to drive expression mostly up, or mostly down, relative to another taxon. This pattern of independent genetic variants with similar effects at similar genes is unlikely under neutral expectations (Orr 1998). It thus serves as a suggestive signature of changes in selective pressure on the pathway between lineages. At this point, rigorous evolutionary conclusions require additional follow-up, including sequence-based tests to distinguish between positive and relaxed selection as the driver of expression divergence. A number of studies in yeast have made this link (Fraser et al. 2012; Martin et al. 2012; Roop et al. 2016); in metazoans, it remains an open question whether results from expression-based *cis*-regulatory pathway tests can be validated with sequence-based molecular-evolution inference (though see Mack et al. 2023).

In the current work, we set out to harness the diversity among mouse lineages in pluripotent stem cell expression programs, to model the integration of expression- and sequence-based tests for selection in multi-gene pathways. We focused on the mouse *Mus musculus castaneus (M. m. castaneus)*. This subspecies is endemic to southeast Asia and diverged 0.5-1 MYA from other *Mus musculus* (Chapman and Ruddle 1972; Sangster et al. 1993). Previous surveys have established divergence between *M. m. castaneus* and laboratory strains in terms of gene expression (Fraser et al. 2011; Xiong et al. 2014; Chappell et al. 2017; Tkatchenko et al. 2019; Chou et al. 2022) and phenotype (Johnson et al. 1997; Beamer et al. 1999; Johnson et al. 2006; Yu et al. 2007; Koturbash et al. 2011; French et al. 2015; Omura et al. 2015; Chappell et al. 2017; Hsiao et al. 2020; Chou et al. 2022), including a particularly avid differentiation phenotype by *M. m. castaneus* stem cells (Ortmann et al. 2020). Our goal was to use stem cell transcriptomes to identify pathways subject to directional *cis*-regulatory change between *M. m. castaneus* and laboratory mice. We earmarked one pathway hit, a set of translation genes at which *M. m. castaneus* alleles acted in *cis* to drive uniquely high expression, for independent validation analyses with sequence data. At these loci, population-genomic tests revealed signals of unique evolution in *M. m. castaneus* in the regions upstream of genes. Together, our results establish the utility of an expression-based molecular-evolution strategy with sequence-based follow-up, and they shed new light on evolutionary and regulatory mechanisms of an ancient divergence in mouse stem cell expression programs.

## Materials and Methods

### RNA-seq data sources

For our initial screen for directional *cis*-regulatory variation in pathways, we used transcriptional profiles of CAST/EiJ male x 129/SvImJ female F1 hybrid embryonic stem cells (NCBI [National Center for Biotechnology Information] accession GSE60738, samples SRR1557132, SRR1557133, SRR1557112, and SRR1557123; Marks et al. 2015). For validation and follow-up we used additional transcriptional profiles from reciprocal crosses of CAST/EiJ x C57BL/6J hybrid pluripotent stem cells (NCBI accession GSE90516, samples SRR5054337-5054348 and SRR5054353-5054364; Werner et al. 2017); homozygous embryonic stem cells from a panel of *M. musculus* subspecies (EBI [European Bioinformatics Institute; Sarkans et al. 2018] accession E-MTAB-7730 [Skelly et al. 2020]); and in-house cultures of the embryonic stem cell line 129Cas, from a male blastocyst from a cross between CAST/EiJ male x 129/SvImJ female, and homozygous induced pluripotent stem cells.

For validation of the role of candidate transcription factors in translation regulation in stem cells, we analyzed published transcriptional data from embryonic stem cells with *Ctr9* knocked down by shRNA (Ruan et al. 2023; NCBI accession GSE219206, samples SRR22493339, SRR22493340, SRR22493341, and SRR22493342) and E8.5 embryos knocked out for *Ehmt2* (Auclair et al. 2016; NCBI accession GSE71500, samples SRR2133432, SRR2133433, SRR2133434, and SRR2133435), respectively.

### RNA-seq read mapping

For analysis of CAST/EiJ x 129/SvImJ (Marks et al. 2015) and CAST/EiJ x C57BL/6J (Werner et al. 2017) hybrid stem cell transcriptomes, we downloaded raw reads and mapped with the STAR aligner (Dobin et al. 2013) to a concatenated genome file containing chromosomal sequences from both parent strains (CAST/EiJ and either 129/SvImJ or C57BL/6J). Data from our in-house cultured hybrid stem cells (see below) were mapped in the same manner.

For analysis of transcriptomes of homozygous CAST/EiJ and C57BL/6J stem cells cultured in-house (see below), we mapped the raw reads to the corresponding reference genome for the respective strain. For analysis of homozygous stem cells from (Skelly et al. 2020) (genotypes C57BL/6J, A/J, 129S1/SvImJ, NZO/HILtJ, NOD/ShiLtJ, WSB/EiJ, CAST/EiJ, and PWD/PhJ), we downloaded raw reads from EBI’s ArrayExpress database (Sarkans et al. 2018) and mapped these to the respective reference genome. For validation of the role of candidate transcription factors in translation regulation in stem cells, we mapped RNA-seq from *Ctr9* knockdown embryonic stem cells (NCBI accession GSE219206; Ruan et al. 2023) and their corresponding shRNA control samples to the 129/SvImJ genome.

### RNA-seq normalization and quantification

Reads that mapped ambiguously to more than one locus were discarded. Read counts were generated during the STAR alignment step using the ‘--quantMode GeneCounts’ option. For each data set in turn, normalized (TPM, transcripts per million) counts were generated by dividing read counts per gene by transcript length (using annotations from the Ensembl database, build 102) and then dividing by library size. Normalized read counts for hybrids and homozygous strains are reported in Table S1 and S2, respectively.

### Hybrid RNA-seq mapping quality control

To eliminate potential artifacts from allele-specific mapping errors in hybrid RNA-seq analyses, we performed a simulated RNA-seq experiment using the Polyester package in R (Frazee et al. 2015) as follows. For the CAST x 129 hybrid stem cell transcriptome from (Marks et al. 2015), we generated two replicates of simulated reads from the hybrid genome with ∼200 reads for each annotated transcript. These simulated reads were then mapped back to the genome with STAR as above. For a given gene with called orthologs in the CAST and 129 genomes, for each allele in turn we tabulated the ratio between the number of successfully mapped simulated reads and the number of simulated reads that went into the mapping, as a report of the extent of artifact-free mapping. We converted each such ratio to a percentage; we then took the absolute value of the difference between the ratio for the 129 and CAST allele as a report of the difference in mapping fidelity between them. We filtered out genes for which the latter parameter exceeded 5%, a total of 1,852 genes of 10,380 initial homologs in the data from (Marks et al. 2015) (Table S3). We repeated this procedure for CAST x BL6 hybrid stem cell transcriptomes from (Werner et al. 2017), resulting in exclusion of 2,886 genes out of an initial 8,887 homologs expressed in the RNA-seq data (Table S3).

### *In silico* screen for directional *cis*-regulatory variation

For our initial screen of polygenic, directional cis-regulatory variation in pathways, we harnessed profiles from CAST/EiJ male x 129/SvImJ female F1 hybrid pluripotent stem cells (Marks et al. 2015). We generated a list of one-to-one orthologous genes (Table S4 between CAST/EiJ and 129/SvImJ from the Ensembl database (build 102) using biomaRt (Durinck et al. 2009). At a given gene, we tested for differential expression between the CAST/EiJ and 129/SvImJ alleles using the reads mapping to each as input into DESeq2 (Love et al. 2014). We eliminated from further analysis genes with fewer than 10 total reads across all samples. Differential expression results are reported in Table S5.

For each gene with differential allele-specific expression at adjusted *p* < 0.05, we assigned a quantitative sign statistic *s*, equal to the log_2_(129 allele expression/CAST allele expression). We assigned all genes without differential allele-specific expression to have *s* = 0. We downloaded gene annotations in Gene Ontology ‘biological process’ terms from the AmiGO database (Carbon et al. 2009). We eliminated from further testing all terms containing fewer than 10 genes with significant differential allele-specific expression. For a given remaining term containing *n* genes, we summed the *s* values across the component genes to yield a summary statistic *S_true_*. To evaluate significance by resampling, we randomly sampled *n* genes from the total set with expression data and summed their *s* values, generating a resampled summary statistic *S_resample_*. We carried out this calculation 10,000 times and used as a two-sided *p*-value the proportion of resamples in which |*S_resample_*| ≥ |*S_true_*|. We corrected for multiple testing with the Benjamini-Hochberg method. *p-*values are reported in Table S6. All further expression and molecular-evolution analyses focused on genes in the top-scoring term, GO:0006412, translation.

### Induced pluripotent stem cell derivation and culture

In-house stem cell expression profiling experiments used material as follows. A 129S6/SvEv embryonic stem cell line was obtained from Millipore Sigma, (Catalog no. SCR012, Burlington, MA, USA). The hybrid embryonic stem cell line was 129Cas, from a male blastocyst from a cross between CAST/EiJ male x 129/SvImJ female, a kind gift from Joost Gribnau. To establish C57BL/6J and CAST/EiJ induced pluripotent stem cell lines, we used mouse embryonic fibroblasts E13.5 obtained from Jackson Laboratories (Bar Harbor, ME, USA) as input into a stem cell derivation protocol as previously described (Terzic et al. 2016). Briefly, we used octamer-binding transcription factor-4 (Oct4), sex-determining region y-box 2 (Sox2), and Kruppel-like factor-4 (Klf4) as reprogramming factors, introduced using pMXs retroviral vectors.

Stem cells were cultured on irradiated mouse embryonic fibroblasts (R and D Systems, Minneapolis, MN, USA) in miPSC medium: knockout DMEM with 4.5 g/ L d-glucose (Gibco, Grand Island, NY, USA), 10% knockout serum replacement (KSR) (Gibco), 10% fetal bovine serum (FBS) (HyClone, Logan, UT, USA), 1× MEM nonessential amino acids (MEM NEAA) (Gibco), 1× GlutaMAX (Gibco), 0.1 mM 2-mercaptoethanol (BME) (Life Technologies, Grand Island, NY, USA), and 0.02% ESGRO-LIF (Millipore, Billerica, MA, USA). Cells were incubated at 37°C in 5% CO_2_.

### RNA isolation and sequencing

RNA was extracted from undifferentiated pluripotent stem cell cultures (two replicates of CAST, four replicates each of 129 and 129Cas) following feeder depletion using the RNAqueous™-Micro Total RNA Isolation Kit (Thermo Fisher Scientific) and on-column DNase treatment (QIAGEN, Hilden, Germany). RNA samples were processed into mRNA libraries and sequenced on an Illumina NovaSeq 6000 Sequencing System, yielding ∼20M paired-end 150 bp reads per sample. Read-mapping was as in RNA-seq normalization and quantification, above.

### Population genomic analysis

We downloaded resequencing data from wild populations of *M. m. domesticus* (from France, Germany and Iran), *M. m. musculus* (from Afghanistan, Czech Republic and Kazakhstan), and *M. m. castaneus* (from northwest India) (Harr et al. 2016). VCF files were used to make two haploid pseudogenomes from each individual mouse in the context of the C57BL/6J reference genome (GRCm38) using the ‘consensus’ command from bcftools software (Danecek et al. 2021), one incorporating the alternate allele at each heterozygous site using the default options, and the other incorporating the reference allele at each heterozygous site using the ‘--haplotype R’ option. We used each pseudogenome separately as input into downstream analyses as described next.

Sequences upstream of the transcription start site for each gene were extracted utilizing the pybedtools Python package (Quinlan and Hall 2010; Dale et al. 2011). In order to assess only a single transcript per gene, gffread (Pertea and Pertea 2020) was used to extract coding sequences (CDS) from each annotated transcript. We retained for analysis only those CDSs that contained an in-frame start and stop codon, signifying a valid open reading frame (ORF). For genes with multiple transcripts containing a valid ORF, the longest transcript was selected.

In a public resource of chromatin immunoprecipitation sequencing data sets (Kolmykov et al. 2021), we tabulated binding sites for 680 transcriptional regulators across the mouse genome. For each regulator, we identified binding sites that were within 50kb upstream of a genic transcription start site (TSS). We analyzed these sites in a comparison of the pseudogenomes from each *M. m. domesticus* or *M. m. musculus* population in turn against the *M. m. castaneus* population as follows. For a given gene and regulator, using the complete set of pseudogenomes for which heterozygote positions had been assigned as the reference allele, we tabulated the number of nucleotide positions in the regulator’s binding sites within 50kb of the TSS that were polymorphic across *M. m. castaneus* and/or the respective sister subspecies population, P_u_. We repeated this calculation using the pseudogenomes in which the heterozygote positions had been assigned as the alternate allele, and we took the mean of the two P_u_ values as a final estimate of the polymorphism of the regulator’s bound site. We used an analogous pipeline to count positions in the regulator’s binding sites that were divergent between *M. m. castaneus* and the respective sister subspecies population and fixed within each, D_u_. Next, we identified all nucleotide positions within 50kb of the TSS that did not fall into binding sites for the regulator, and we calculated P_u_ and D_u_ as above. We carried out these calculations for all genes and all populations. We then used the resulting values, per regulator, as input into a two-factor ANOVA, testing for an interaction between binding site identity and gene membership in the translation GO term, using the populations of *M. m. domesticus* and *M. m. musculus* as replicates. We applied this approach to each transcriptional regulator and corrected for multiple testing with the Benjamini-Hochberg method (Benjamini and Hochberg 1995). Significant ANOVA results are reported in Table 1; full results are in Table S7. Multiple sequence alignment was edited for display using Jalview software (Waterhouse et al. 2009). Binding site data for the top four hits from this analysis with an odds ratio > 1 are reported in Table S8. To check whether the D_u_/P_u_ enrichment in binding sites of our top-scoring regulator, Ctr9, was driven by patterns of divergence or polymorphism, we repeated the two-factor ANOVA test as above but used as input D_u_ by itself and, separately, P_u_ by itself (normalized by binding site length), rather than the ratio between them.

## Results

### A screen for directional *cis*-regulatory change in mouse stem cell pathways

As a testbed for analyses of pathway *cis*-regulatory change, we used transcriptional profiles (Marks et al. 2015) of embryonic stem cells of an F1 hybrid background from a cross between two homozygous mouse strains: a male *M. m. castaneus* (CAST/EiJ, hereafter CAST), and a female of the 129/SvImJ laboratory strain genotype (hereafter 129), which is of admixed origin (Frazer et al. 2007; Yang et al. 2007; Yang et al. 2011). In the F1, because the two subspecies’ alleles of a given gene are in the same nucleus, any difference in allele-specific expression between them can be attributed to genetic variation acting in *cis*, *i.e.,* not mediated through a soluble factor (Wittkopp et al. 2004). We implemented a pipeline of allele-specific read-mapping taking account of the potential for mapping artifacts (see Methods); and we retained for analysis all genes exhibiting significant expression divergence between the species’ alleles (962 genes at a 0.05 *p*-value threshold). We tabulated the directional effect for each gene—whether the CAST allele was more highly expressed than the laboratory-strain allele, or *vice versa*—using the log_2_-transformed fold-change between expression of the 129 allele and CAST allele. Then, to formulate our test, we used as pathways groups of genes of common function, each comprised of a Gene Ontology biological process term. For each such group, we quantified the agreement in the direction of allelic expression differences between species across the gene members. We evaluated significance based on resampling (Table S6). Of the complete survey results, one pathway showed significant signal: a cohort of genes from the GO term for translation, at which the CAST allele was upregulated relative to that of 129 in the hybrid at a 2-fold excess of genes relative to those with higher expression from the 129 allele (Table S6, Figure 1A, and Figure S1A). This represented a potential case in which selective pressures on regulation of the respective loci had changed between the *M. m. castaneus* and laboratory-strain lineages.

**Figure 1.**
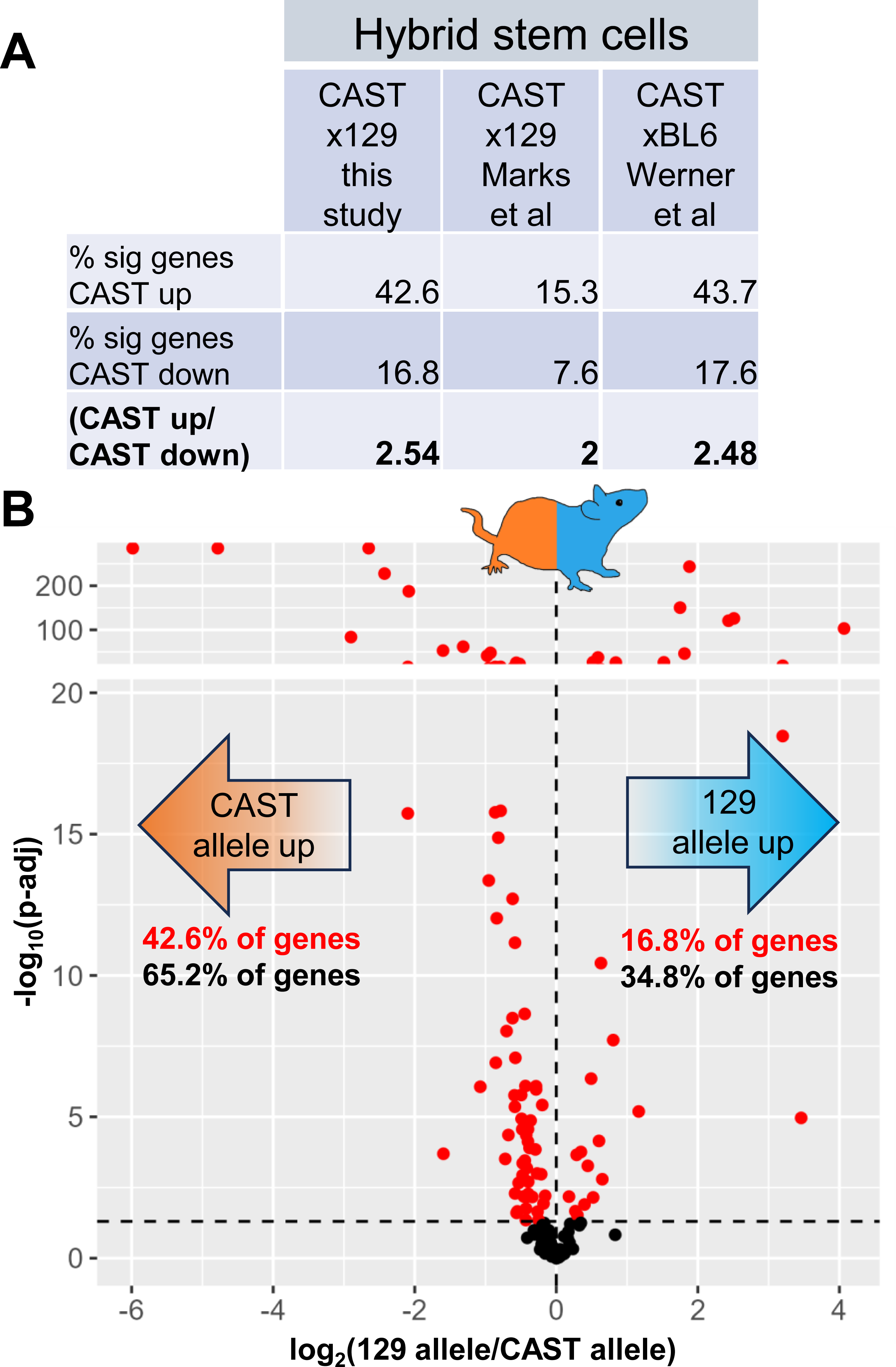
Directional *cis*-regulatory variation in translation gene expression in stem cells between mouse subspecies. **(A)** Results from analyses of differential allele-specific expression across genes from the Gene Ontology term GO:0006412, translation, in stem cells of F1 hybrids between CAST and either C57BL/6 (from Werner et al. 2017) or 129 (Marks et al. 2015 or this study) as indicated. The first and second rows report the percentage of translation genes in which the CAST allele is expressed higher or lower, respectively, than the allele of the other parent, and the third row reports the ratio of these quantities. **(B)** Each point reports allele-specific expression of a translation gene in CASTx129 hybrid stem cells cultured in this study: the *x*-axis reports the log_2_ ratio of expression of the respective strain alleles, and the *y*-axis reports the log_10_ of the significance of the difference (p-adj, adjusted *p*-value). Point colors report significance of differential allele-specific expression (red, adjusted *p* < 0.05; black, adjusted *p* > 0.05). Red and black text inlays report the percentage of translation genes where expression of the respective parental allele is higher with and without filtering for significance, respectively.

### *M. m. castaneus cis*-regulatory alleles drive high expression of translation genes in stem cells

As a first verification of the trend for *cis*-regulatory alleles from *M. m. castaneus* driving high expression of translation genes, we repeated the culture and sequencing of CAST x 129 hybrid stem cells. The results confirmed the robust directional imbalance in allele-specific expression among translation genes, with the CAST allele expressed more highly across the set 2.54-fold more often than the 129 allele (Figure 1A-B). Similarly, we analyzed the transcriptome of hybrid stem cells from a cross between CAST and the admixed C57BL/6J laboratory strain (hereafter BL6; Werner et al. 2017), and observed a 2.48-fold imbalance favoring high expression by the CAST allele among translation genes (Figure 1A and Figure S1B). Together, these data establish that *M. m. castaneus cis*-regulatory alleles at translation genes encode a unique activating program relative to those encoded by 129 and BL6 alleles, in stem cells.

### Homozygous *M. m. castaneus* stem cells are distinguished by high expression of translation genes

We expected that, if expression divergence between *M. m. castaneus* and other lineages at translation genes had been important for fitness in the organismal and ecological context, it would be apparent in the context of homozygous strains, which integrate the effects of genetic factors acting both in *cis* and in *trans* (Signor and Nuzhdin 2018). Consistent with this prediction, translation genes were more highly expressed in CAST homozygous stem cells relative to those of 129 and BL6, in a published data resource (Skelly et al. 2020) and in our own culture and sequencing (Figure 2A-B and Figure S1C-D). The trend persisted in comparisons between CAST homozygous stem cells and those of other mouse lineages (Figure 2A and Figure S1E-I). Interestingly, the tendency for high expression of translation genes by CAST in homozygous stem cells was of larger magnitude than we had noted in our analyses of *cis*-regulatory variation in hybrids (Figure 1A), indicating that the latter was reinforced by a stronger effect of divergence attributable to *trans-*acting regulators. Given these signatures of directional *cis*-and *trans*-acting variation between *Mus* subspecies, we considered the translation gene cohort to be an informative target for mechanistic and molecular-evolution follow-up.

**Figure 2.**
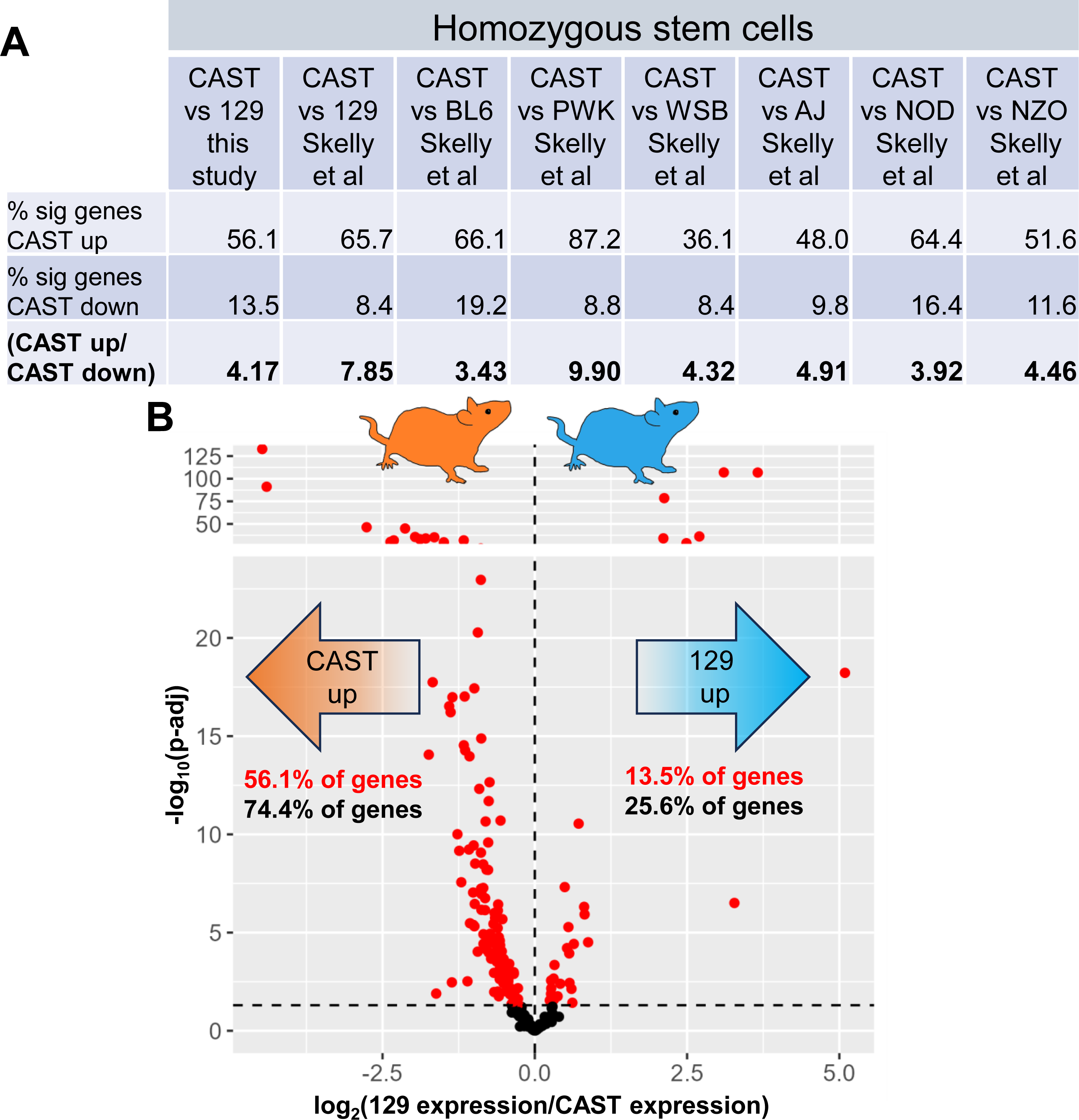
Directional *trans*-regulatory variation in translation gene expression in stem cells between mouse subspecies. **(A)** Data are as in Figure 1A except that each column reports results from comparisons between homozygous CAST and homozygous 129 (this study or Skelly et al. 2020), C57BL/6J, A/J, NZO/HILtJ, NOD/ShiLtJ, WSB/EiJ, CAST/EiJ, or PWD/PhJ (Skelly et al. 2020) as indicated. **(B)** Data are as in Figure 1B except that each point reports expression of a translation gene in a comparison between homozygous CAST and 129 stem.

### Unique alleles in *M. m. castaneus* at regulatory loci of translation genes

To dissect regulatory and evolutionary mechanisms of the expression divergence in translation genes between *M. m. castaneus* and other lineages, we focused on sequence changes that could serve as candidate determinants of *cis*-regulatory variation, upstream of coding regions. For this purpose we developed a screening pipeline for single-nucleotide variants from wild-caught mice (Harr et al. 2016), which we classified on the basis of upstream binding sites by transcriptional regulators (Kolmykov et al. 2021). For the sites bound by a given regulator, we evaluated divergence between *M. m. castaneus* and its relatives, normalized by intra-subspecies polymorphism, across the genes of our translation cohort, in light of the classic interpretation of this metric as a hallmark of positive selection. Results revealed robust differences in normalized divergence between the translation genes and a genomic null in the binding sites of each of seven regulators (Table 1). In four of these top-scoring cases, binding sites at translation genes were enriched for normalized divergence, indicative of an excess of derived alleles that distinguished *M. m. castaneus* from *M. m. domesticus* and *M. m. musculus* (Table 1). All four such screen hits were well-studied regulators of cell identity and differentiation (Ctr9, Rfx6, Hoxa11, and Ehmt2). An additional three regulators emerging from our population-genomic screen had significantly low normalized divergence between *M. m. castaneus* and its relatives in binding sites upstream of translation genes, reflecting especially limited variation between subspecies at these positions and/or relaxed constraint within them (Table 1). We conclude that evolutionarily interpretable changes, most notably spikes of alleles unique to *M. m. castaneus* suggestive of positive selection, can be resolved at *cis*-regulatory sites in translation genes. And these divergent variants emerge across the binding sites for multiple regulators, as expected if evolutionary rewiring in this system involved a complex network of inputs.

### Signatures of evolutionary and regulatory impact of Ctr9 site variants at translation genes

As a test case for deeper insights into mouse subspecies variation in the translation pathway, we focused on Ctr9, a core member of the Paf1/RNA polymerase II complex whose binding sites were top-scoring in terms of normalized divergence between subspecies at our focal genes (Table 1). The signal manifested in each comparison between *M. m. castaneus* and a given *M. musculus* relative (Figure 3A-B). Analyses of the components of our normalized metric made clear that the latter result was driven by elevated variation between subspecies per se: divergence between *M. m. castaneus* and its relatives was 1.7-fold higher in Ctr9 sites at translation genes than at control loci (Figure 3C), whereas polymorphism within populations at these sites was almost on par with that of controls (Figure 3D). Given these patterns of unique alleles in *M. m. castaneus* at Ctr9 sites at translation genes, we considered them as particularly likely to contribute to the *cis*-controlled *M. m. castaneus* expression program that we had noted in this pathway in stem cells (Figure 1). Consistent with such a function, Ctr9 sites with sequence divergence between *M. m. castaneus* and its relatives coincided with divergent expression by the *M. m. castaneus* allele in hybrid stem cells, among translation genes (Figure 3E). Furthermore, our inference of Ctr9 as an activator for the translation pathway was borne out by expression profiles of *Ctr9* knockdown in stem cells of laboratory mice (Ruan et al. 2023), which established a marked directional influence of Ctr9 on translation genes (Figure 3F). Together, these data highlight Ctr9 as a case in which binding sites for a stem cell transcriptional regulator have evolved uniquely at translation genes in *M. m. castaneus*, representing candidate drivers of non-neutral expression divergence in this system.

**Figure 3.**
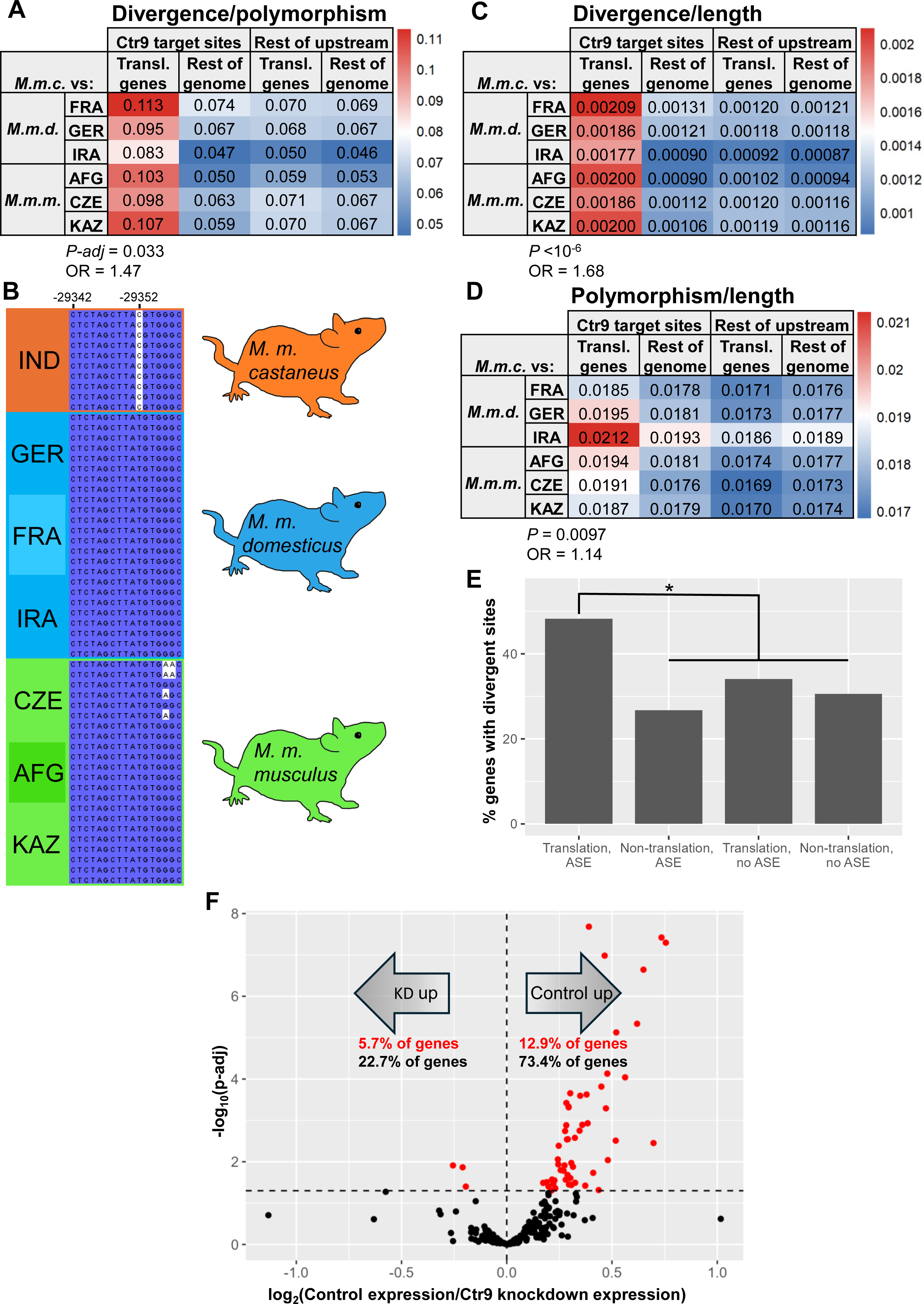
Signatures of species divergence and regulatory function of Ctr9 at translation genes. **(A)** In a given row, each colored cell reports normalized divergence, the ratio of inter-subspecies divergence to within-subspecies polymorphism, at Ctr9 binding sites upstream of genes from the Gene Ontology term GO:0006412, translation, or all other genes of the genome, in a comparison between *M. m. castaneus* and the indicated population of a sister taxon. FRA, France; GER, Germany; IRA, Iran; AFG, Afghanistan; CZE, Czech Republic; KAZ, Kazakhstan. Cells are colored based on relative values. Values at the bottom report the results of an ANOVA testing for differences in normalized divergence between translation genes and controls (see Table 1 and Table S7). *P-adj*, screen *p*-value after correction for multiple testing; OR, odds ratio. **(B)** A portion of a Ctr9 binding footprint upstream of the translation gene Rps29 as an example of mouse subspecies variation. The *x*-axis reports position relative to the transcription start site. Each row reports the sequence of one mouse individual; at heterozygote loci, only the allele representing the alternate to the reference is shown, although our analysis methods accounted for both (see Methods). **(C)** Data are as in A, except that each value reports the number of sites divergent between *M. m. castaneus* and the indicated sister population in the region of interest, standardized by the length of the latter. **(D)** Data are as in A, except that each value reports the number of sites polymorphic within *M. m. castaneus* and/or the indicated sister population in the region of interest, standardized by the length of the latter. **(E)** Shown is an analysis of mouse subspecies divergence variation at Ctr9 binding sites, in genes stratified by membership in the translation gene cohort and allele-specific expression differences between CAST and 129 alleles in F1 hybrid stem cells (ASE; Marks et al. 2015; Table S5). For a given bar, the *y*-axis reports the percentage of genes in the indicated category containing variants in Ctr9 binding sites divergent between *M. m. castaneus* and one or more sister taxon (Table S8). *, Fisher’s exact test *p* = 0.02. **(F)** Data and symbols are as in Figure 1B except that each point reports a comparison of expression of one translation gene between *Ctr9* knockdown and shRNA control mouse stem cells (Ruan et al. 2023), and red and black text inlays report the percentage of translation genes where expression was higher in the indicated genotype with and without filtering for significance, respectively.

## Discussion

Empirical molecular evolution tests, once developed, must be validated to establish their rigor and utility for the field. A key means toward this end is to integrate results from an emergent test strategy with those of more classic tools, when they complement each other in support of an evolutionary inference. In the current work, we have detected sequence divergence between mouse subspecies in a cohort of translation genes that also exhibits directional, polygenic *cis*- and *trans*-acting regulatory variation in stem cells. Against a backdrop of other case studies focused on *cis*-regulatory change as detected in transcriptomes (Bullard et al. 2010; Fraser et al. 2010; Fraser et al. 2011; Martin et al. 2012; York et al. 2018; Agoglia et al. 2021; Gokhman et al. 2021; Mack et al. 2023; Wang et al. 2024), our results represent a proof of concept for sequence-based validation of this approach in mammals.

Our findings in mouse subspecies echo reports of *cis*-regulatory change in the translation machinery of other organismal systems, from yeasts (Tanay et al. 2005; Hogues et al. 2008; Li and Fay 2017; Sorrells et al. 2018) to animal ancestors distributed over deep time (Brown et al. 2008). This literature leaves open the question of what ecological forces might drive divergence of translation gene expression, and the phenotypes that would mediate such effects. Under a model in which ribosomal protein dosage governs the readiness of a cell to divide (Polymenis and Aramayo 2015), adaptive regulatory variation in translation pathways may often reflect species-unique logic of cell growth decisions.

In metazoan stem cells in particular, translation plays a critical role in differentiation. Inducing (Easley et al. 2010) or compromising (Khajuria et al. 2018) stem cell translation can drive qualitative differences in differentiation behavior. According to current models, high expression of translation genes in stem cells (Sampath et al. 2008) sets up a poised state to enable rapid protein production in their differentiated progeny (Gabut et al. 2020). On the basis of this tight link between translation and differentiation, it is tempting to speculate that the *M. m. castaneus* expression program in translation genes could underlie the uniquely avid differentiation behavior by stem cells of this genotype into definitive endoderm (Ortmann et al. 2020). Such a phenotype could well represent an adaptation, as suggested by the expression-and sequence-based signatures of selection we detect for the divergent regulation at translation genes in *M. m. castaneus*. More broadly, we would expect translation factors to act as part of a complex genetic architecture of stem cell differentiation as it differs between mouse lineages, alongside other variants mapped in this system (Ortmann et al. 2020; Skelly et al. 2020).

Our population-genomic screen highlighted several transcriptional regulators whose binding sites harbor unique alleles in *M. m. castaneus* at translation genes, each of which represents a candidate determinant of the stem cell expression program in this lineage. The case for function of these regulators in stem cells is supported by a deep prior literature from laboratory mice. The top hit from our screen, Ctr9, is an integral member of the Pol II-associating factor 1 complex (Mueller and Jaehning 2002) with a well-characterized role in development and maintaining stem cell identity and pluripotency (Ding et al. 2009; Ruan et al. 2023). Direct binding by Ctr9 to translation genes in stem cells (Rahl et al. 2010; Ding et al. 2020) dovetails with our inference of Ctr9 regulation of this cohort, including its difference between lineages. Given the occupancy of Ctr9 at superenhancers as well as downstream of transcription termination sites in embryonic stem cells (Ding et al. 2020), such elements may ultimately prove to underlie Ctr9 function at translation loci.

Likewise, the methyltransferase Ehmt2, another hit in our screen for unique alleles in *M. m. castaneus* at translation genes, has well-known functions in stem cell maintenance and differentiation (Ikegami et al. 2007; Leitch et al. 2013; Boroviak et al. 2014; Auclair et al. 2016; Kim et al. 2020) and binds directly to translation genes in stem cells (Mozzetta et al. 2014). Indeed, early embryonic knockout of Ehmt2 (Auclair et al. 2016) had a directional impact on expression of translation genes in stem cell transcriptomes (Figure S2). Our screen hit Hox11a, a well-studied developmental regulator (Yamamoto and Kuroiwa 2003; Boulet and Capecchi 2004; Wong et al. 2004; Horvat-Switzer and Thompson 2005; Kherdjemil et al. 2016; Rux et al. 2016; Leclerc et al. 2023), has been characterized in a similar literature, including descriptions of its function in stem cells (Rux et al. 2016; Song et al. 2020; Leclerc et al. 2023). And Rfx6, also a hit in our screen, has been previously implicated in development in a number of animal systems (Smith et al. 2010; Trott et al. 2020; Lu et al. 2021), including in translation gene regulation (Cheng et al. 2019; Lu et al. 2021). The emerging picture from our analyses is thus that Ctr9, Rfx6, Hoxa11, and Ehmt2 may exert their effects on stem cell identity and differentiation in part via regulation of protein synthesis, and that binding sites for these regulators represent candidate players in the mechanism by which evolution has tuned expression of translation factors in *M. m. castaneus*. That said, we expect that our genomic approach affords only partial coverage and power in discovering elements of this mechanism, and that many other contributing regulators likely remain to be identified.

In summary, our discovery of a unique expression program in *M. m. castaneus* stem cells, at translation genes which also harbor divergent binding sites for a suite of regulators, represents an informative molecular-evolution case study. And our results serve as a foundation for future work at the physiological level, to pursue the relevance of translation gene regulation in stem-cell differentiation (Ortmann et al. 2020) and other phenotypes (Johnson et al. 1997; Beamer et al. 1999; Johnson et al. 2006; Yu et al. 2007; Koturbash et al. 2011; French et al. 2015; Chappell et al. 2017; Hsiao et al. 2020; Chou et al. 2022) that distinguish *M. m. castaneus* from the rest of its genus.

## Acknowledgements

The authors thank Kaylee Christensen for assistance with computational resources and Taekyu Kang and Matt Dean for helpful discussions. This work was supported by National Institutes of Health R01NS116992 to DMB, JRD, and RBB, and R01GM120430 to RBB.

## Author Contributions

NMS, RBB, DMB, and JRD conceived of the study and provided direction. NMS processed the sequencing data and carried out all *in silico* analyses. JRD derived the iPSC lines used for sequencing and neuron differentiation. YK performed the cell cultures and RNA isolations. All authors contributed to and reviewed the final manuscript.

## Data Archiving

RNA-seq data generated for this study was deposited in the GEO database (https://www.ncbi.nlm.nih.gov/geo/query/acc.cgi?acc=GSE234761).

## Table captions

**Table 1. Patterns of variation between mouse subspecies in binding sites of transcriptional regulators upstream of translation genes.** Shown are results of analysis of normalized sequence divergence, at the regions upstream of genes, between *M. m. castaneus* on the one hand and *M. m. musculus* and *M. m. domesticus* on the other. Each row reports results of an ANOVA testing for an interaction effect between two factors on the divergence metric: position of sequence variants in binding sites of the indicated transcriptional regulator and gene membership in the translation GO term (see Figure 3A). Normalized divergence was calculated as the number of sites divergent between *M. m. castaneus* and a given *M. musculus* relative, normalized by the number of sites polymorphic within the populations. The second and third columns report the odds ratio (OR) and nominal *p*-value, respectively, and the third reports the *p*-value after Benjamini-Hochberg correction for multiple testing. OR > 1 indicates enrichment of normalized divergence at regulator binding sites relative to the other three categories, and OR < 1 indicates depletion. Full results are shown in Table S7.

## Supplementary figure captions

**Figure S1. Directional *cis*-regulatory variation in translation gene expression in stem cells between mouse subspecies.** Data are as in Figure 1B **(A-B)** or Figure 1C **(C-I)** of the main text except that transcriptomes analyzed were as follows. **(A)** CAST x 129 hybrid stem cells (Marks et al. 2015). **(B)** CAST x BL6 hybrid stem cells (Werner et al. 2017). **(C)** CAST and 129 homozygous stem cells (Skelly et al. 2020). **(D)** CAST and BL6 homozygous stem cells (Skelly et al. 2020). **(E)** CAST and WSB/EiJ homozygous stem cells (Skelly et al. 2020). **(F)** CAST and PWD/PhJ homozygous stem cells (Skelly et al. 2020). **(G)** CAST and A/J homozygous stem cells (Skelly et al. 2020). **(H)** CAST and NOD/ShiLtJ homozygous stem cells (Skelly et al. 2020). **(I)** CAST and NZO/HILtJ homozygous stem cells (Skelly et al. 2020).

**Figure S2. Ehmt2 knockout induces translation genes.** Data are as in Figure 3C of the main text except that transcriptomes analyzed were wild-type and Ehmt2 knockout stem cells (Auclair et al. 2016).

## Supplementary table captions

**Table S1. Allele-specific normalized read counts from RNA-seq of stem cells from hybrids involving CAST.** Each tab reports allele-specific expression measurements in one genotype of hybrid embryonic stem cells. Tab TPM_GSE60738_CAST_M_x_129_F contains results from four replicate transcriptomes of replicates of CAST/EiJ male x 129/SvImJ female F1 hybrid cells from (Marks et al. 2015). Tab TPM_GSE234761_CAST_M_x_129_F contains results from four replicate transcriptomes of 129Cas (CAST/EiJ male x 129/SvImJ female F1 hybrid) cells from this study. Tab TPM_GSE90516_CAST_M_x_BL6_F contains results from 12 replicate transcriptomes of CAST/EiJ male x C57BL/6J female cells (Werner et al. 2017). Tab TPM_GSE90516_CAST_F_x_BL6_M contains results from 11 replicate transcriptomes of CAST/EiJ female x C57BL/6J male cells (Werner et al. 2017). In each tab, the first column reports Ensembl gene accession numbers. The remaining columns are TPM (transcripts per million) values for reads mapped non-ambiguously to either parental allele for each gene (two columns for each biological sample).

**Table S2. Normalized read counts from RNA-seq of stem cells from homozygous strains.** Each tab reports expression measurements from one data source of experimental profiles of cell culture from homozygous mouse strains: pluripotent stem cells (PSCs) of 129 and CAST/EiJ genotypes from this study; embryonic stem cells of eight genotypes from (Skelly et al. 2020); *Ctr9* knockdown in embryonic stem cells (Ruan et al. 2023); and *Ehmt2* knockouts in E8.5 embryos (Auclair et al. 2016). In each tab, the first column reports Ensembl gene accession numbers. The remaining columns report the TPM (transcripts per million) values. Tab TPM_GSE234761_homozyg_CAST_129 contains results from two replicate transcriptomes of CAST PSCs and four replicate transcriptomes of 129 PSCs. Tab TPM_E_MTAB_7730_8_strains contains results from ESC replicate transcriptomes from eight strains (3 replicates CAST, 35 replicates PWD, 3 replicates WSB, 36 replicates BL6, 3 replicates 129, 3 replicates AJ, 22 replicates NOD, 3 replicates NZO). Tab TPM_GSE219206_CTR9_KD contains results from two replicate transcriptomes of control shRNA ESC samples and two replicate transcriptomes of shRNA knockdown of *Ctr9,* Tab TPM_GSE71500_EHMT2_KO contains results from two replicate transcriptomes of wild-type embryos (E8.5) and two replicate transcriptomes of *Ehmt2* knockout embryos (E8.5).

**Table S3. Mapping results from simulated RNA-seq experiments using hybrid genomes.** The first and second tabs report the mapped read counts from simulated RNA-seq of CAST x 129 and CAST x BL6 hybrids, respectively. In a given tab, for each row, the first column lists the Ensembl gene id for the gene analyzed; the second through fourth columns report the mapping percentage (the number of simulated reads mapped, normalized by the total number of simulated reads) for the indicated allele in the indicated simulated replicate; the last column reports the absolute value of the difference of mapping percentages between the CAST allele and that of the indicated strain (129 or BL6).

**Table S4. Accession ID numbers for gene homologs across strains.** Gene and transcript accession numbers used to compare homologous alleles between strains. GRCm38_ensembl_gene_ID, Ensembl gene IDs for the C57BL/6 reference. 129_MGP_gene_ID, 129S1/SvImJ Mouse Genome Project IDs. CAST_MGP_gene_ID, CAST/EiJ Mouse Genome Project IDs. gene_name, gene names from the Mouse Genome Informatics database. GRCm38_ensembl_transcript_ID is the Ensembl transcript ID for the longest in-frame ORF for that gene; empty cells in this column are genes with no valid ORF in the GRCm38 genome using the Ensembl build 102 annotations.

**Table S5. Differential expression test results in hybrid and homozygous stem cells.** Each tab reports results of a genome-wide survey of differential expression between homologous alleles in interspecies hybrid stem cells; between homozygote stem cells of different species genotypes; or between a regulator gene knockout and its isogenic wild-type in stem cell or early embryo samples. Log2FoldChange, log-transformed ratio of normalized expression of the indicated alleles or homozygote samples; Pvalue, Wald test *p*-value; Padj, Benjamini-Hochberg adjusted *p*-value. Tab DESeq_GSE60738_CASTx129_hyb, comparison of CAST vs 129 allele-specific expression in F1 hybrid stem cells from (Marks et al. 2015). Tab DESeq_GSE234761_CASTx129_hyb, comparison of CAST vs 129 allele-specific expression in F1 hybrid stem cells from this study. Tab DESeq_GSE90516_CAST_x_BL6_hyb, comparison of CAST vs BL6 allele-specific expression in F1 hybrid stem cells from (Werner et al. 2017). Tab DESeq_GSE234761_CASTvs129, comparison of CAST vs 129 expression in homozygous stem cells from this study. Tab DESeq_E_MTAB_7730_CASTvs129, comparison of CAST vs 129 expression in homozygous stem cells from (Skelly et al. 2020). Tab DESeq_E_MTAB_7730_CASTvsBL6, comparison of CAST vs BL6 expression in homozygous stem cells from (Skelly et al. 2020). Tab DESeq_E_MTAB_7730_CASTvsWSB, comparison of CAST vs WSB expression in homozygous stem cells from (Skelly et al. 2020). Tab DESeq_E_MTAB_7730_CASTvsPWD, comparison of CAST vs PWD expression in homozygous stem cells from (Skelly et al. 2020). Tab DESeq_E_MTAB_7730_CASTvsAJ, comparison of CAST vs AJ expression in homozygous stem cells from (Skelly et al. 2020). Tab DESeq_E_MTAB_7730_CASTvsNOD, comparison of CAST vs NOD expression in homozygous stem cells from (Skelly et al. 2020). Tab DESeq_E_MTAB_7730_CASTvsNZO, comparison of CAST vs NZO expression in homozygous stem cells from (Skelly et al. 2020). Tab DESeq_GSE219206_Ctr9_KD, comparison of control shRNA vs *Ctr9* knockdown expression in stem cells from (Ruan et al. 2023). Tab DESeq_GSE71500_EHMT2_KO, comparison of wildtype vs *Ehmt2* knockout expression in E8.5 embryos from (Auclair et al. 2016).

**Table S6. A screen for directional allele-specific expression in CAST x 129 hybrid embryonic stem cells.** Each row reports results of a statistical test for directional allele-specific expression variation in CAST/EiJ male x 129/SvImJ female F1 hybrid embryonic stem cells (Marks et al. 2015) in the indicated Gene Ontology term. total_number_of_genes, number of genes in the term with analyzable expression measurements; sum_sign_statistic, sum across genes of the term of log_2_(129 allele expression/CAST allele expression); resampling_P_val, resampling-based significance of the enrichment for high absolute value of the sign statistic in the term; Adjusted_P, *p*-value after Benjamini-Hochberg correction for multiple testing.

**Table S7. Patterns of variation between mouse subspecies in binding sites of transcriptional regulators upstream of translation genes.** Data are as in Table 1 of the main text, except that results from all tested regulators are shown.

