## Supplementary figures and images for "Stem cell transcriptional profiles from mouse subspecies reveal *cis*-regulatory evolution at translation genes"

### Figure S1

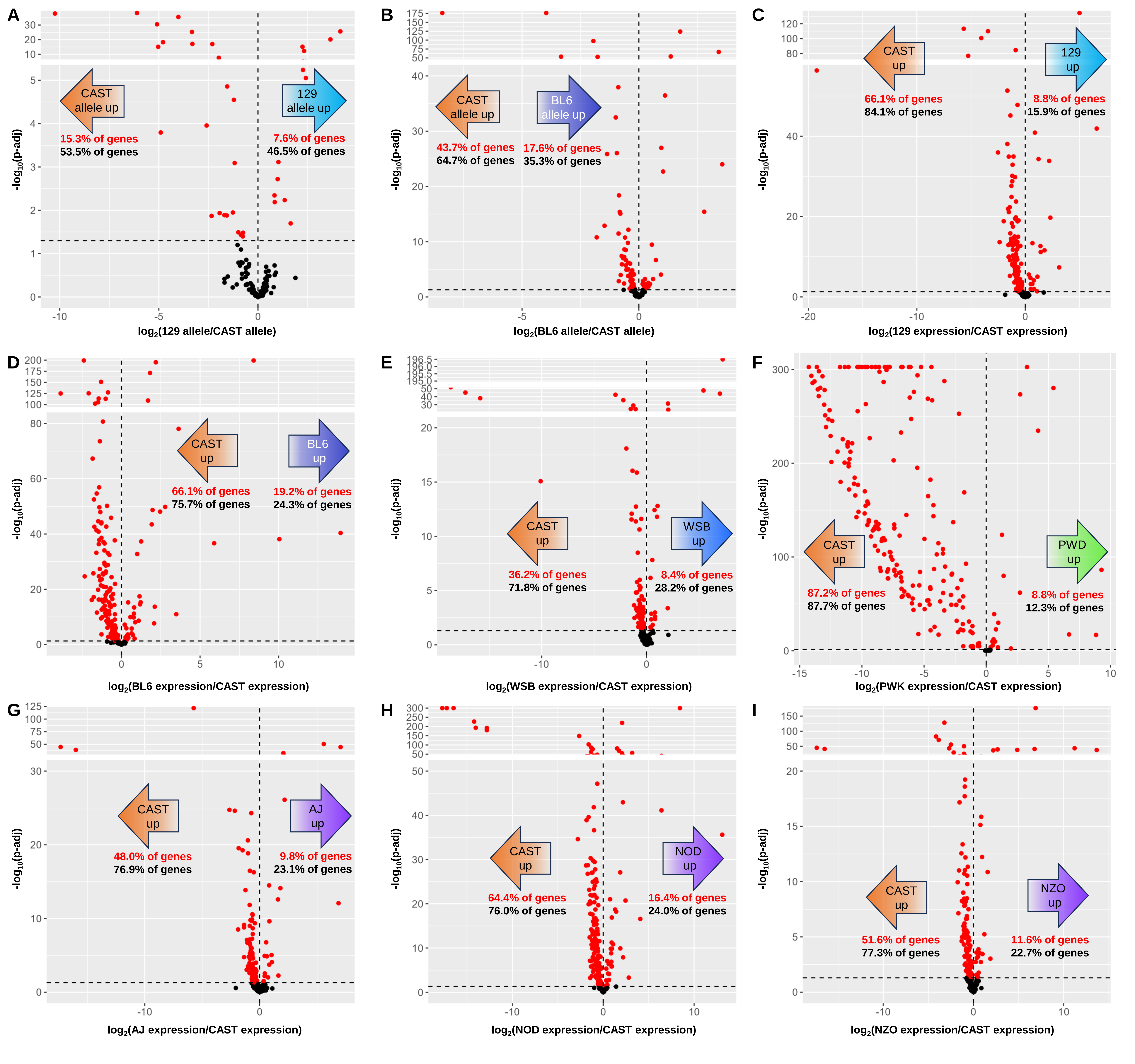

### Figure S2

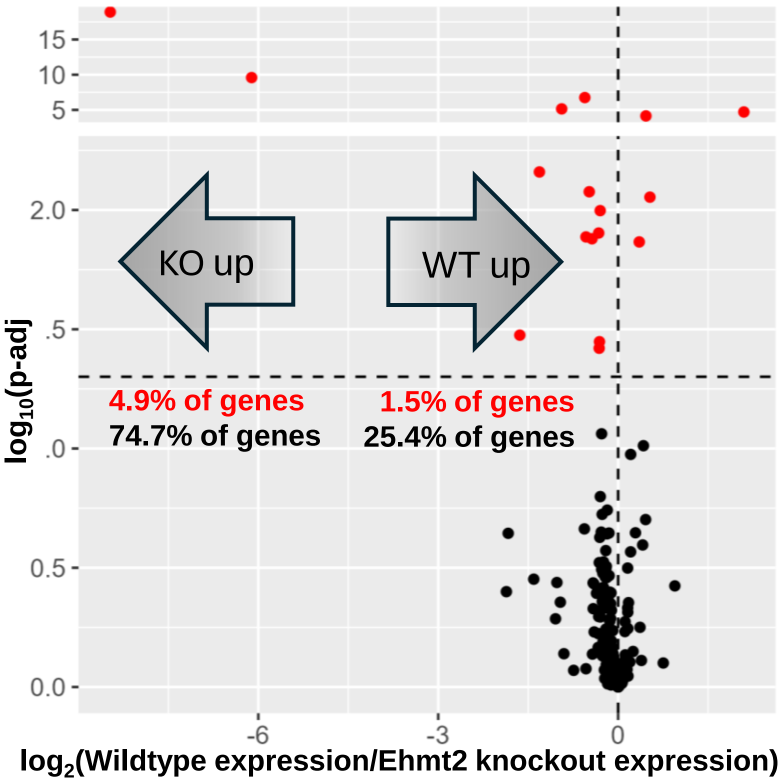
